# Spatial Transcriptomics Reveals Region-Specific Remodeling in Vein Grafts After Peripheral Arterial Bypass

**DOI:** 10.64898/2026.06.08.731009

**Authors:** Keisuke Kamada, Hong Niu, Shinsuke Kikuchi, Nobuyoshi Azuma, Gale L. Tang

## Abstract

**Background:** Vein graft failure due to intimal hyperplasia and maladaptive remodeling remains a major limitation of peripheral bypass surgery. Although vascular remodeling is recognized as a multilayered process, layer-specific molecular mechanisms that distinguish adaptive from negative remodeling remain incompletely understood. We aimed to investigate the vascular microenvironment of patent and stenotic grafts using spatial transcriptomics.

**Methods:** Vein specimens were obtained from three patients undergoing revision surgery. For each patient, a matched set of three samples was collected: unused saphenous vein (Denovo), normally healed vein graft (Non-stenosed), and stenosed vein graft (Stenosed) (n = 3 patients). GeoMx Digital Spatial Profiling with the Human Whole Transcriptome Atlas was used to map gene expression across intima, medial, and adventitial layers. Differential expression, gene ontology, spatial deconvolution, and immunohistochemistry were integrated for analysis.

**Results:** Non-stenosed and Stenosed grafts shared transcriptional features distinct from Denovo veins, particularly in pathways related to cell proliferation. Non-stenosed grafts showed increased expression of *CDKN1A* across all vascular layers, whereas Stenosed grafts exhibited enhanced mitogen-activated protein kinase (MAPK) pathway activity, reduced *DUSP1*-mediated regulation, and increased macrophage accumulation. ECM remodeling showed layer-specific organization, with *VCAN* and *ACAN* enriched in the intima and *DCN* in the adventitia, while Stenosed grafts demonstrated a trend toward collagen-dominant remodeling. Cell deconvolution suggested a shift toward vascular smooth muscle cell (VSMC)-dominant architecture after arterialization, with modest enrichment of synthetic VSMC signatures in stenotic regions.

**Conclusions:** Vein graft stenosis appears to be associated with layer-specific alterations in cell cycle regulation, inflammatory signaling, extracellular matrix remodeling, and VSMC phenotype. Spatial transcriptomic analysis reveals molecular heterogeneity not captured by bulk approaches and provides preliminary insight into graft remodeling. These findings may inform future studies to improve long-term graft patency.

## Introduction

Peripheral arterial disease affecting the lower extremities is a major contributor to limb ischemia, morbidity, and mortality worldwide.^1, 2^ In patients with chronic limb-threatening ischemia, surgical bypass using autologous vein grafts remains a cornerstone of revascularization—particularly when endovascular strategies are insufficient or anatomically unsuitable.^3, 4^ Among conduits, the great saphenous vein (GSV) is preferred for its favorable long-term patency compared to prosthetic alternatives in peripheral bypasses, but graft failure remains problematic. In the distal bypass setting, graft failure occurs in 20–30% of vein grafts within 1 to 2 years postoperatively due to intimal hyperplasia (IH) or negative remodeling.^5, 6^ This incidence has persisted despite advances in surgical technique, perioperative management, and surveillance imaging. Understanding the mechanisms of IH in peripheral vein grafts remains incomplete, particularly regarding layer-specific processes and interlayer interactions. Vein graft remodeling is a multifactorial process involving vascular smooth muscle cell (VSMC) migration and proliferation, extracellular matrix (ECM) reorganization, inflammatory signaling, and hemodynamic stress. Despite extensive investigation of these mechanisms, the spatial organization of molecular programs within human peripheral vein grafts remains poorly defined.^7–12^

While molecular and translational studies have advanced our understanding of vein graft remodeling, spatially resolved analyses of human peripheral vein grafts remain limited. Consequently, layer-specific transcriptional programs distinguishing adaptive from stenotic remodeling remain poorly defined. Previous studies have suggested that VSMCs and adventitial cells isolated from human GSVs exhibit distinct responses to proliferative stimuli and differences in gene expression profiles, implying that each vascular layer may play unique and independent roles in the development of vein graft stenosis.^9, 13^ Bulk RNA sequencing of peripheral graft tissues can capture average expression changes but fails to resolve spatial heterogeneity or layer-specific phenomena. Meanwhile, animal graft models, though valuable, do not fully replicate the human vascular and inflammatory environment in the lower limb—including variable shear stress, metabolic milieu, and comorbid influences.

Spatial transcriptomics represents a promising modality to bridge this gap: it allows gene expression profiling within defined histologic regions while preserving spatial architecture, enabling dissection of intimal, medial, and adventitial compartments. Recent advances in spatial transcriptomics have led to its application in cardiovascular research, including studies of myocardial disorders and atherosclerotic plaques.^14, 15^ However, despite these developments, spatially resolved transcriptomic analyses of human peripheral bypass vein grafts remain scarce, leaving layer-specific mechanisms of graft adaptation and failure largely unexplored.

In this study, we applied the GeoMx^®^ Digital Spatial Profiler to define the spatial transcriptomic architecture of clinically relevant human vein grafts. Importantly, we directly compared unused GSV segments (Denovo), normally healed graft segments (Non-stenosed), and stenotic graft segments (Stenosed) obtained from the same patients undergoing revision surgery, enabling analysis across distinct stages of graft adaptation and failure. Histology-guided annotation of intimal, medial, and adventitial compartments allowed layer-resolved interrogation of molecular programs within preserved tissue architecture. Furthermore, integration of spatial transcriptomic profiling with cell deconvolution and pathway-level analyses provided a systems-level framework for understanding graft remodeling. Through this approach, we sought to identify coordinated molecular signatures and cellular phenotypes that distinguish adaptive healing from pathological stenosis. These findings provide mechanistic insight into vein graft failure and offer a translational basis for developing targeted strategies to improve long-term graft patency.

## Methods

Please see the Major Resources Table in the Supplemental Materials.

## Data Availability

Main data generated or analyzed in this study are included in this article. Further inquiries can be directed to the corresponding author.

## Study Approval

This study was approved by the Institutional Review Board of Asahikawa Medical University (Approved number: 19152). Research transparency was ensured by notifying participants through an opt-out approach.

## Human Sample Collection

This study included three cases of vein graft stenosis following peripheral arterial bypass surgery, in which graft revision procedures were performed at Asahikawa Medical University Hospital between January and December 2023. As shown in the protocol flowchart (**Fig. 1A**), the stenosed vein graft was resected with an adequate margin of adjacent non-stenosed portion and replaced with an unused segment of the GSV from ipsilateral or contralateral limb. The excised graft was then divided into a normally healed segment (**Non-stenosed**; n=3) and a stenosed segment (**Stenosed**; n=3). In addition, a leftover portion of de novo GSV (**Denovo**; n=3) was collected. All samples were fixed in 10% neutral buffered formalin (NBF) for at least 24 hours and subsequently embedded in paraffin blocks. To optimize slide usage, 4–5 graft samples were embedded in a single block (multi-tissue block). Sections were cut at a thickness of 5 μm and stored at 4 °C until further analysis.

**Figure 1.**
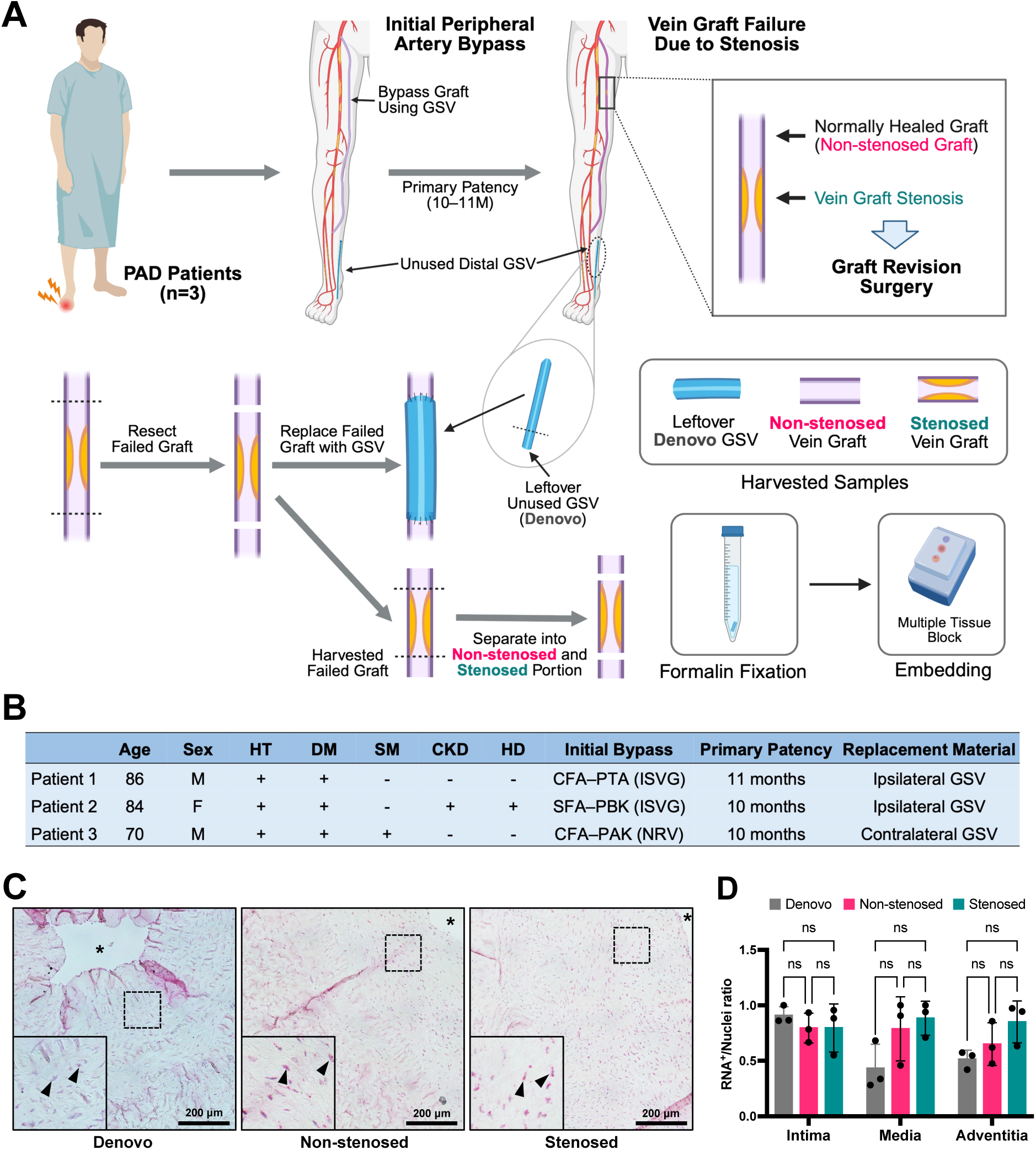
Patient background and RNA quality assessment of vein graft tissues embedded in paraffin blocks. **A.** Flowchart of graft revision procedures and sample collection. Vein graft samples were obtained from patients undergoing revision surgery for graft failure. Failed grafts were divided into non-stenosed and stenosed potions, and unused GSV segments were also collected. **B.** Patient clinical background and graft primary patency period. **C.** Representative RNAscope image captured by brightfield microscope. RNA signals were indicated as pink punctate dots (arrow head). Lumen area was marked as *. Scale bar = 200 *μ*m. **D.** Quantification of RNA-positive signal relative to nuclei across graft types (Denovo; n=3, Non-stenosed; n=3, Stenosed; n=3). *GSV*, great saphenous vein; *HT*, hypertension; *DM*, diabetes mellitus; *SM*, smoking history; *CKD*, chronic kidney disease; *HD*, hemodialysis. *CFA*, common femoral artery; *SFA*, superficial femoral artery; *PTA*, posterior tibial artery; *PAK*, popliteal above knee artery; *PBK*, popliteal below knee artery; *ISVG*, in-situ vein graft; *NRV*, non-reversed vein

## Morphological Assessment

Morphological evaluation was performed using hematoxylin and eosin (H&E) and Movat’s pentachrome staining on consecutive sections prepared for DSP analysis. Movat’s staining was performed using a modified protocol to improve elastin staining reliability, enabling simultaneous visualization of elastic fibers, collagen, proteoglycans, and muscle components, as previously described.^16^

## RNAscope In-situ Hybridization (ISH)

RNAscope^TM^ ISH was performed on formalin-fixed paraffin-embedded (FFPE) vein graft sections according to the manufacturer’s protocol (Advanced Cell Diagnostics, Newark, CA). Sections were baked at 60 °C, deparaffinized, and rehydrated, followed by endogenous peroxidase blocking with 3% hydrogen peroxide. Target retrieval was performed at 100 °C, and protease digestion was conducted at 40 °C.

Slides were hybridized with the positive control probe Hs-Ppib (ACD #313901, Advanced Cell Diagnostics) at 40 °C, followed by sequential signal amplification. Detection was achieved using Fast Red chromogen. Sections were counterstained with hematoxylin. RNA quality was assessed by quantification of RNA-positive signal relative to nuclei across graft types, confirming adequate RNA preservation for subsequent DSP analysis.

## Slide preparation for Spatial Transcriptomic Study

Slide preparation for spatial transcriptomic analysis was performed according to the manufacturer’s protocol for the GeoMx DSP. All equipment and containers were treated with RNase-inactivating solution prior to use. FFPE sections were baked at 60 °C, deparaffinized, rehydrated, and subjected to antigen retrieval in Tris/ethylenediaminetetraacetic acid buffer (pH 9.0), followed by Proteinase K treatment. After fixation in 10% NBF, slides were hybridized overnight at 37 °C with spatial RNA profiling probes.

After stringent washes, slides were blocked and incubated with fluorescent morphological marker antibodies, including rabbit anti-CD45 (1:100, #1317, Cell Signaling, Danver, MA) and mouse anti–α-smooth muscle actin (α-SMA, 1:100, #53-9760, Invitrogen, Waltham, MA), followed by fluorophore-conjugated secondary antibodies (Goat anti-Rabbit IgG-AF647, 1:100, #111-605-144, Jackson ImmunoResearch, West Grove, PA). Nuclei were counterstained with Syto83 (1:10, #S11364, Invitrogen). Slides were then scanned using the GeoMx DSP instrument to visualize morphological marker staining, enabling layer-specific annotation of regions of interest (ROIs) for downstream analysis.

## ROI selection and Library Preparation

ROIs were selected using the GeoMx DSP instrument with reference to H&E-stained serial sections. For each graft sample, 2–4 ROIs were randomly selected from each histological layer, with the requirement that each ROI contained at least 100 nuclei. In total, 84 ROIs were obtained across the three layers and vein groups from three patients and collected into a 96-well plate.

Sequencing libraries were prepared according to the manufacturer’s instructions and sequenced on an Illumina NovaSeq 6000 S2 platform with a 2 × 50 bp paired-end configuration. For comparative analyses, gene expression was compared across graft types (Denovo, Non-stenosed, and Stenosed) within each vascular layer (intima, media, and adventitia), with each ROI treated as an independent replicate.

## Data Processing

FASTQ files generated from sequencing were processed using the GeoMx NGS Pipeline to generate digital count conversion files, which were subsequently imported into the GeoMx DSP Analysis Suite (version 3.1.4). Quality control was performed by applying the following criteria: a raw read threshold of 1,000, ≥80% of reads aligned, and ≥50% sequencing saturation. After additional filtering (5%) and Q3 normalization, a total of 80 ROIs were retained for downstream analysis, comprising Denovo (n = 24: intima, n=6; media, n=9; adventitia, n=9), Non-stenosed (n = 27: intima, n=9; media, n=9; adventitia, n=9), and Stenosed (n = 29: intima, n=10; media, n=10; adventitia, n=9).

## Differential Expression and Gene Ontology Analysis

Differentially expressed gene (DEG) analysis was performed using the GeoMx DSP Analysis Suite, with cut-off thresholds set at |log_2_ fold change| > 1 and –log_10_ p-value > 1.3. Based on the DEG dataset, gene ontology (GO) enrichment analysis was conducted using Metascape. Among the enriched terms, those related to cell cycle and cell proliferation, as well as inflammation-associated pathways, were specifically extracted for further interpretation.

## Cell Deconvolution and Gene Set Enrichment Analysis

Cell deconvolution analysis was performed using the SpatialDecon R package, with a publicly available single-cell RNA sequencing dataset (Tabula Sapiens – Vasculature) obtained from CZ CELLxGENE Discover as the reference profile matrix (https://cellxgene.cziscience.com).

To further explore VSMC and macrophage phenotypes, single-sample Gene Set Enrichment Analysis (ssGSEA) was conducted using the ssGSEA R package. For VSMC phenotypes, the following gene sets were applied: contractile VSMC (*MYH11*, *ACTA2*, *TAGLN*, *CNN1*, *SMTN*) and synthetic VSMC (*FN1*, *SPP1*, *MMP2*, *MMP9*, *VCAN*).^17–19^ For macrophage phenotypes, the following gene sets were used: M1 macrophages (*IL1B*, *IL6*, *CCL2*, *STAT1*, *IRF1*) and M2 macrophages (*MRC1*, *CD163*, *TGFB1*, *MSR1*),^20, 21^ with normalization to CD68 expression.

## Immunofluorescence Staining

To evaluate ECM components and macrophage phenotypes, immunofluorescence staining was performed using the following primary antibodies: anti-mouse versican (Working concentration 1:500, MA5-27638, Thermo Fisher Scientific, Waltham, MA), anti-mouse aggrecan (1:500, MCA1454G, Bio-Rad, Hercules, CA), anti-rabbit decorin (1:50, PA5-95830, Thermo Fisher Scientific), anti-mouse CD86 (1:100, 942-MSM1-P1ABX, Thermo Fisher Scientific), and anti-rabbit CD206 (1:2000, ab64693, Abcam, Cambridge, UK). The following secondary antibodies were used: Alexa Fluor 488 goat anti-rabbit (1:300, A32731TR, Invitrogen), Alexa Fluor 647 goat anti-rabbit (1:300, A21245, Invitrogen), and Alexa Fluor 488 goat anti-mouse (1:300, A11001, Invitrogen). Sections were counterstained with 4′,6-diamidino-2-phenylindole (DAPI; 1 μg/mL, D5942, Sigma-Aldrich, St. Louis, MO). Negative control staining (omission of the primary antibody) was also performed (data not shown). For each patient, two to four randomly selected fields (×20 magnification) were imaged using a confocal microscope (Nikon AX/AX R with NSPARC, Nikon). ImageJ/Fiji software (National Institutes of Health) was used to quantify primary antibody staining and DAPI intensity ratio. The mean value of 2–4 images per patient was calculated, and these averages were used for analysis (n = 3 patients).

## Statistical Analysis

All statistical analyses were performed using Prism 10 (GraphPad Software, Inc., Boston, MA) and R Studio (R version 4.4.3). For DSP data analysis, each ROI was treated as an independent replicate, whereas the patient number (biological replicate) was used for quantification of immunohistochemical staining. Comparisons between two independent groups were conducted using Student’s t-test, and comparisons among multiple groups were analyzed by two-way ANOVA followed by Tukey’s multiple comparisons test. Data are presented as mean ± standard deviation (SD). Levels of significance are indicated as follows: \**P* < 0.05, \*\**P* < 0.01, \*\*\**P* < 0.001, and \*\*\*\**P* < 0.0001.

## Results

### Patient Background and Morphological Assessment

Human vein graft samples were obtained from patients who underwent graft revision surgery following initial peripheral bypass grafting (**Fig. 1A**). All patients had a history of hypertension and diabetes. One patient had end stage kidney disease requiring hemodialysis. The primary patency period was approximately 10–11 months (**Fig. 1B**).

Movat’s pentachrome staining was performed to evaluate morphological differences between Non-stenosed and Stenosed vein grafts. (**Fig. S1A**). The fold of increase in neointimal thickness was greater in Stenosed than in Non-stenosed (\**P*<0.05, **Fig. S1B**). The intimal and medial thickness ratios, normalized to Denovo, were significantly higher in Stenosed than in Non-stenosed grafts (**Fig. S1C**).

To assess RNA quality and confirm the suitability of the samples for DSP analysis, RNAscope ISH was conducted. The ratio of RNA-positive signal to nuclei tended to be lower in the media and adventitia of Denovo; however, no significant differences were observed among the groups (**Fig. 1C, 1D**). These results indicated that all samples were suitable for DSP experiments.

### Differentially Expressed Genes Between Graft Types for Each Layer

The workflow of spatial transcriptomic profiling from FFPE blocks is shown in **Figure S2A**. ROIs were selected based on immunostaining for CD45 and α-SMA, with reference to H&E staining on consecutive sections (**Fig. 2A**, **Fig. S2B**).

**Figure 2.**
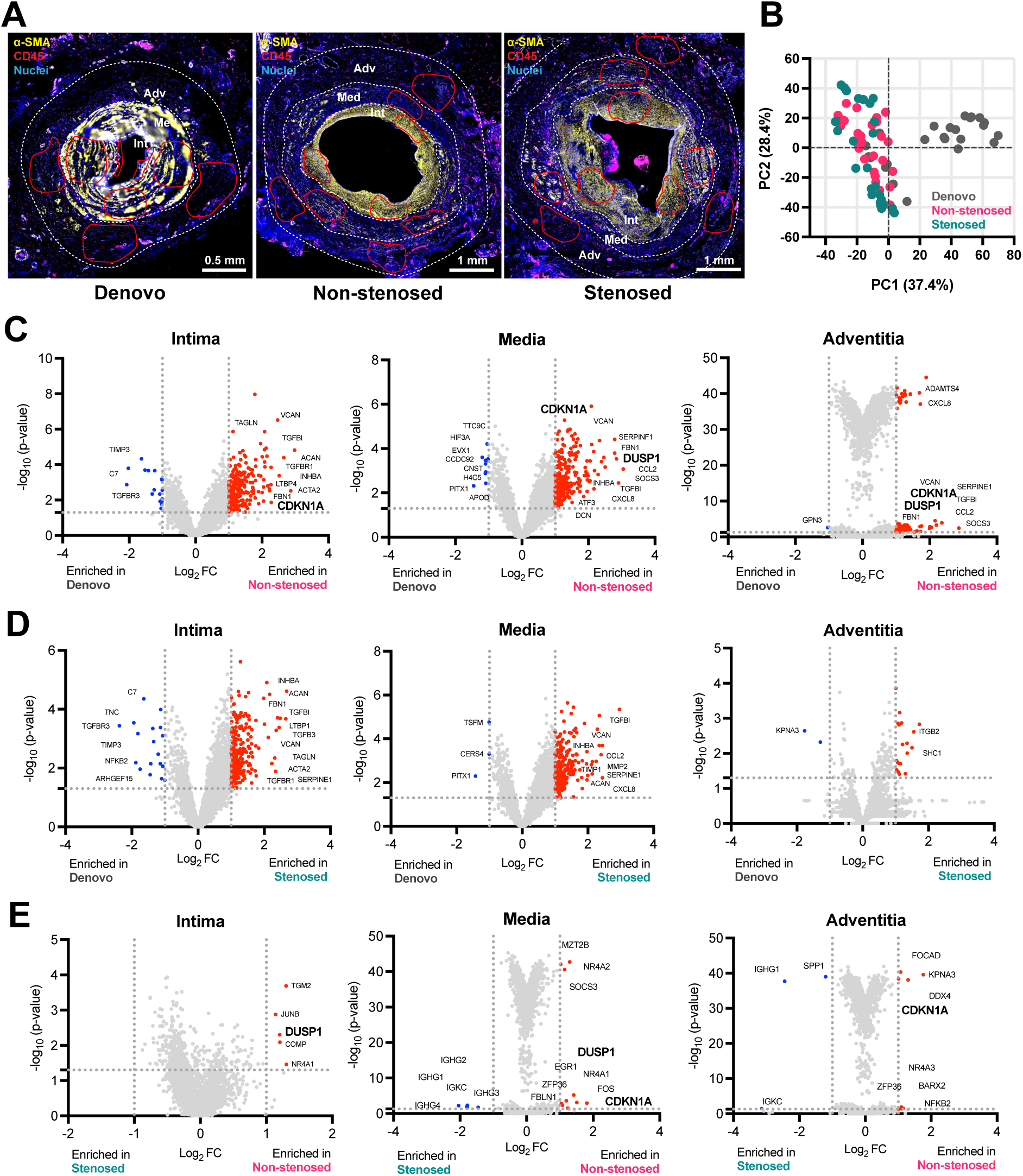
Spatial transcriptomic profiling and representative differentially expressed genes (DEGs) in intima, media, and adventitia. A. ROI selection based on CD45 and α-SMA immunostaining. The regions were identified using the corresponding H&E images. B. Principal component analysis of Denovo (ROI, n = 24), Non-stenosed (ROI, n = 27), and Stenosed (ROI, n = 29). Volcano plots showing DEGs for each comparison: C. Denovo vs Non-stenosed, D. Denovo vs. Stenosed, and E. Stenosed vs. Non-stenosed.

Principal component analysis demonstrated that Non-stenosed and Stenosed shared similar transcriptional features, which were distinct from those of Denovo (**Fig. 2B**).

DEG analysis revealed characteristic differences among the graft types. Representative genes associated with key biological processes, including cell cycle, cell proliferation, inflammation, and extracellular matrix remodeling, are highlighted in the figure. In the comparison between Denovo and Non-stenosed, the cell cycle inhibitor gene *CDKN1A* was enriched across the intima to adventitia, and the mitogen-activated protein kinase (MAPK) regulator *DUSP1* was enriched in the media and adventitia (**Fig. 2C**). In contrast, Stenosed did not show similar enrichment of these genes relative to Denovo (**Fig. 2D**). Furthermore, when comparing Non-stenosed with Stenosed, *CDKN1A* was more highly expressed in the media and adventitia, and *DUSP1* was enriched in the intima and media of Non-stenosed (**Fig. 2E**).

### Regulation of Cell Cycle and Proliferation in Non-stenosed Graft

We next examined genes and pathways related to cell cycle regulation and cellular proliferation. *CDKN1A* was significantly upregulated in the media of Non-stenosed grafts compared with both Denovo and Stenosed grafts (vs. Denovo, \*\**P* <0.01; vs. Stenosed, \**P*<0.05). *LMNA*, a nuclear structural protein that also plays a role in cell cycle regulation, was enriched in the intima, media, and adventitia of Non-stenosed grafts relative to Denovo. In addition, the cell proliferation inhibitor *IGFBP3* was highly expressed in the intima of Non-stenosed compared with Denovo and Stenosed (**Fig. 3A**). Similar trends were also observed for *IGFBP7*, *THBS1*, and *INHBA* (**Fig. S3**).

**Figure 3.**
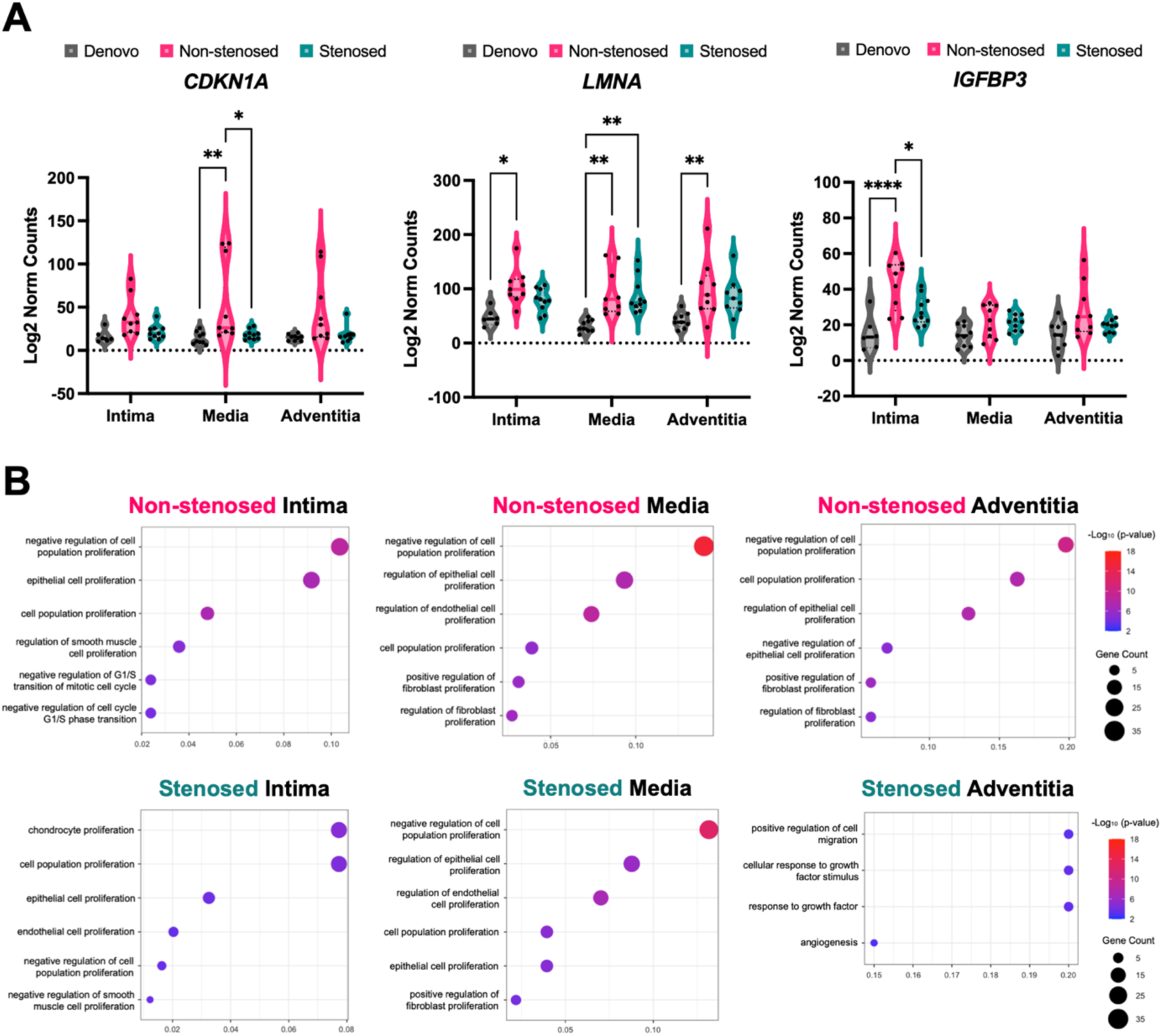
Distinct regulation of cell cycle and proliferation between Non-stenosed and Stenosed grafts. A. Expression of representative genes associated with cell cycle or cell proliferation: *CDKN1A* (cell cycle inhibitor), *LMNA* (a nuclear structural protein that also plays a role in cell cycle regulation), and *IGFBP3* (cell proliferation regulator). Denovo (ROI, n = 24: intima, n=6; media, n=9; adventitia, n=9), Non-stenosed (ROI, n = 27: intima, n=9; media, n=9; adventitia, n=9), and Stenosed (ROI, n = 29: intima, n=10; media, n=10; adventitia, n=9). B. GO analysis based on DEG datasets, highlighting cell cycle- or cell proliferation-associated GO terms.

GO analysis was then performed using the data from DEG analysis. The top six enriched GO terms for each graft type and histological layer were selected and visualized. GO terms related to cell cycle regulation and cell proliferation were extracted by graft type and histological layer.

Non-stenosed was enriched for pathways associated with negative regulation of cell population proliferation across the intima, media, and adventitia, whereas Stenosed grafts exhibited enrichment for this pathway only in the media (**Fig. 3B**).

### Differences in Inflammatory Regulation Among Graft Types

We assessed inflammation related genes and macrophage associated genes across different groups. *DUSP1*, a regulator of the MAPK pathway, was significantly upregulated in the intima and media of Non-stenosed grafts compared with Stenosed grafts (Non-stenosed intima, \**P*<0.05; Non-stenosed media, \*\**P*<0.01). The inflammation-associated regulator *ZFP36* was enriched in the adventitia of Non-stenosed relative to both Denovo and Stenosed. In contrast, the general macrophage marker *CD68* was significantly higher in the adventitia of Stenosed than in Denovo and Non-stenosed (\*\**P*<0.01, **Fig. 4A**). Other inflammation-related markers, including *IL6*, *MRC1*, and *HMGB1*, showed no significant differences among the groups (**Fig. S4A**). Macrophage polarization was further assessed by CD86 (M1) and CD206 (M2) co-staining. Although CD86 tended to be higher in the adventitia of Stenosed grafts compared with Denovo and Non-stenosed grafts, the difference did not reach statistical significance; similarly, CD206 showed no significant variation (**Fig. S4B, S4C**). In addition, ssGSEA was conducted to assess macrophage phenotypes based on the gene sets. While no significant differences were observed in the intima, both Non-stenosed and Stenosed showed higher M1 macrophage enrichment scores in the media and adventitia compared with Denovo (\*\**P*<0.01). No significant enrichment was observed for M2 macrophages (**Fig. S4D**).

**Figure 4.**
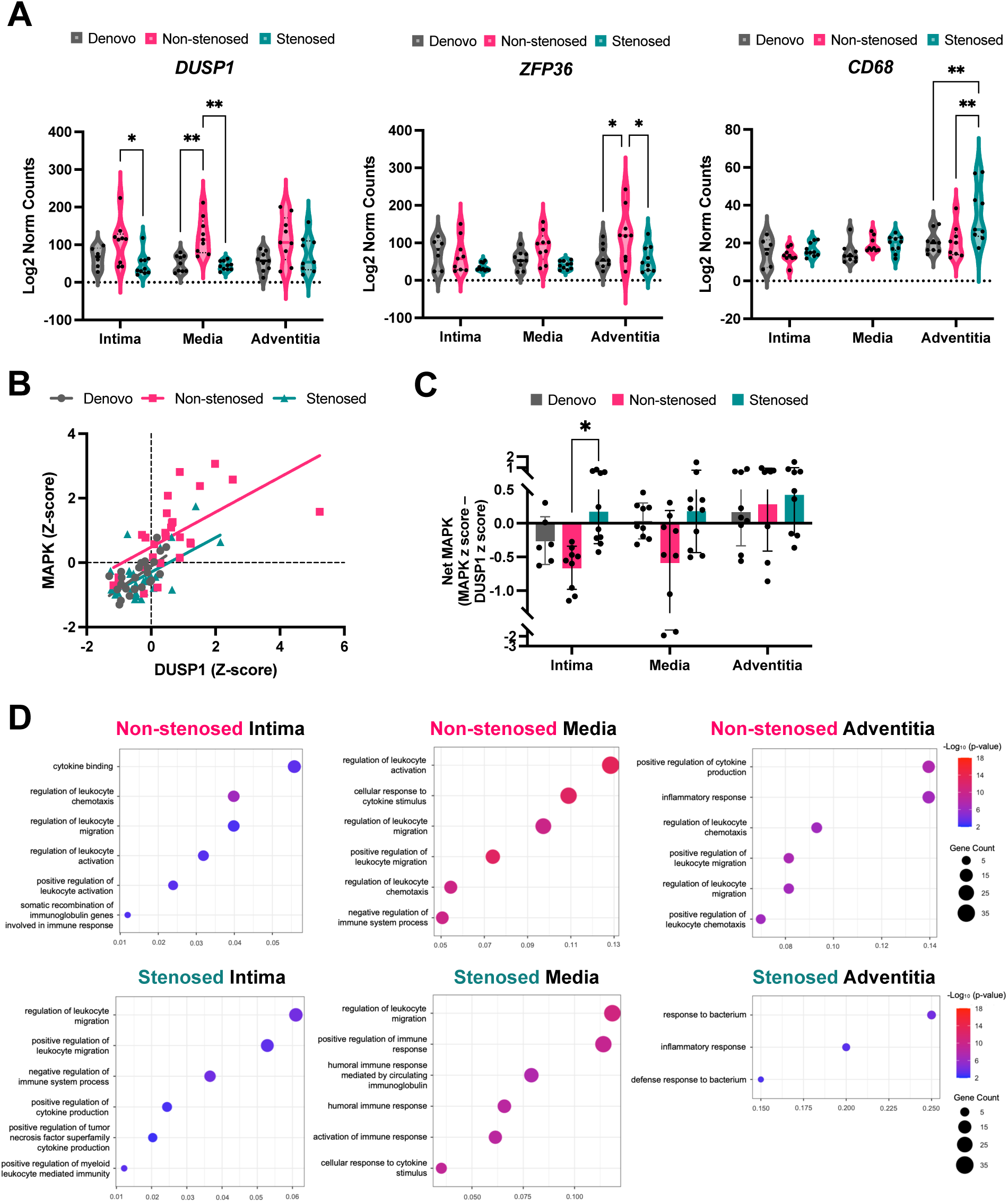
Inflammatory regulation among graft types. **A.** Expression of inflammation-associated markers: *DUSP1* (MAPK regulator), *ZFP36* (inflammation-associated regulator), and *CD68* (general macrophage marker). **B.** A positive correlation was observed between DUSP1 z-score and MAPK z-score. **C.** Net MAPK score (MAPK z-score – DUSP1 z-score). The Net MAPK score was significantly higher in the intima of Stenosed compared with Non-stenosed. **D.** GO terms related to inflammatory regulation tended to be more enriched in Non-stenosed than in Stenosed across all layers. Denovo (ROI, n = 24: intima, n=6; media, n=9; adventitia, n=9), Non-stenosed (ROI, n = 27: intima, n=9; media, n=9; adventitia, n=9), and Stenosed (ROI, n = 29: intima, n=10; media, n=10; adventitia, n=9). *MAPK*, mitogen-activated protein kinase

For further assessment of the relationship between *DUSP1* and MAPK pathway activity, a MAPK z-score was calculated based on the expression of *FOS*, *JUN*, *EGR1*, *DUSP5*, *DUSP6*, *IL1B*, *IL6*, and *CXCL10*.^22–24^ The expression of *DUSP1* was likewise converted to a z-score, enabling comparison with the MAPK z-score. A positive correlation was observed between these two parameters (**Fig. 4B**). To further delineate MAPK activity, the Net MAPK score was derived by subtracting the *DUSP1* z-score from the MAPK z-score. The Net MAPK score was significantly higher in the intima of Stenosed compared with Non-stenosed (\**P*<0.05, **Fig. 4C**). GO terms associated with inflammatory regulation tended to be more frequently enriched in Non-stenosed than in Stenosed across all layers (**Fig. 4D**).

### Regulation of ECM Gene Expression and Protein Distribution Across Graft Types

ECM remodeling is a critical component of graft adaptation. To investigate ECM dynamics, we focused on three key proteoglycans — versican (*VCAN*), aggrecan (*ACAN*), and decorin (*DCN*) — and assessed their expression across layers and graft types. *VCAN* and *ACAN* exhibited similar patterns, with both being significantly upregulated in the intima of Non-stenosed and Stenosed compared with Denovo. *DCN* expression was significantly higher in the adventitia of Non-stenosed and Stenosed than in their intima (\*\**P* < 0.01, **Fig. 5A**). We next evaluated other ECM-related genes. *COL1A1* was significantly elevated in the intima and adventitia of Stenosed relative to Denovo (\**P*<0.05). While *MMP9* showed no significant difference, *MMP2* expression was consistently higher in the adventitia compared with the intima across all graft types (Denovo and Non-stenosed graft, \*\**P*<0.01; Stenosed graft, \*\*\*\**P*<0.0001). In addition, *TIMP1* (\*\**P*<0.01) and *ADAMTS4* (\**P*<0.05) were enriched in the media and adventitia of Non-stenosed, respectively (**Fig. S5**).

**Figure 5.**
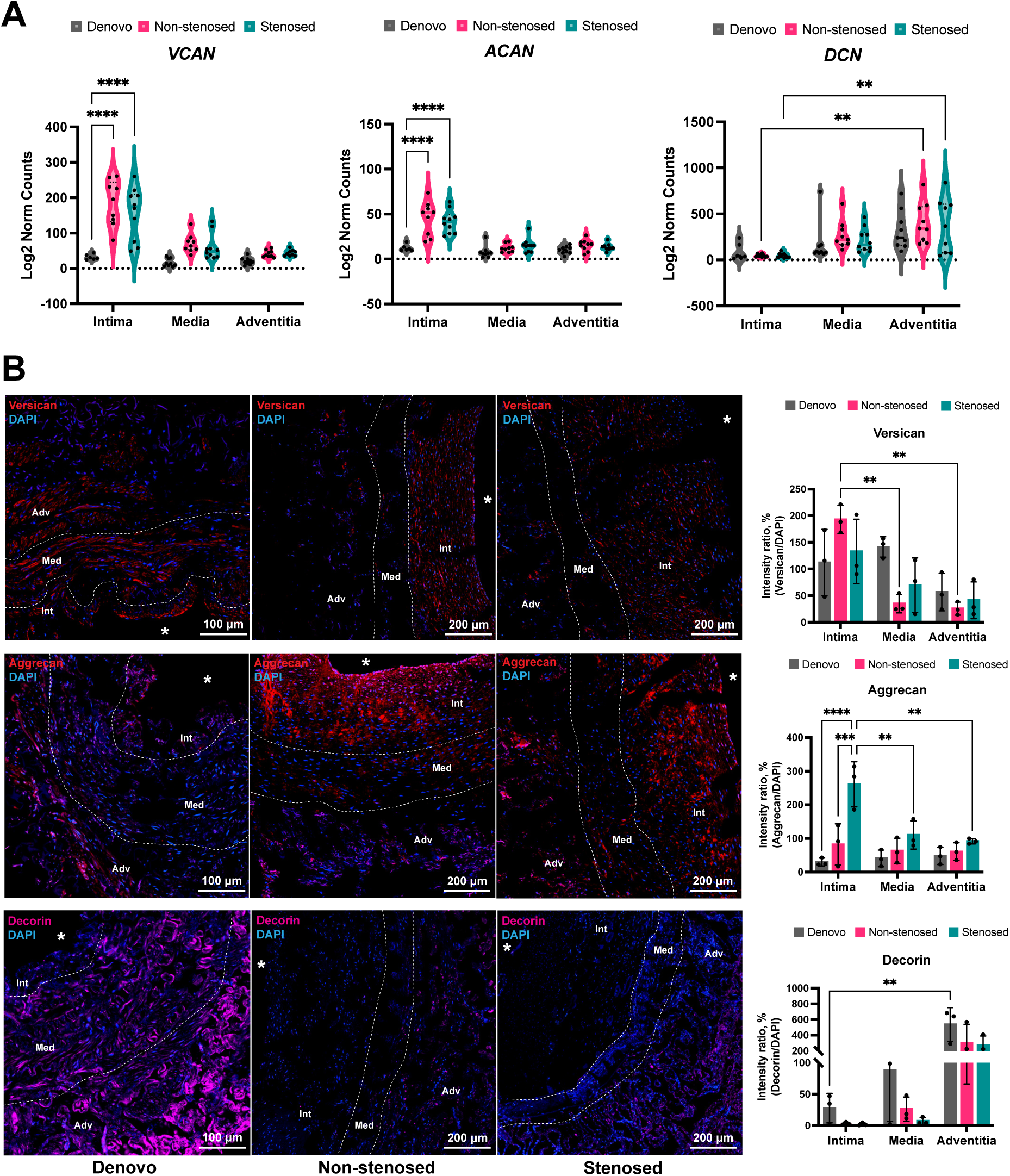
Regulation of ECM gene expression and protein distribution. **A.** Expression of proteoglycan genes: *VCAN* (Versican), *ACAN* (Aggrecan), and DCN (Decorin). *VCAN* and *ACAN* were upregulated in the intima of Non-stenosed and Stenosed compared with Denovo, whereas *DCN* was higher in the adventitia than in the intima in both groups. Denovo (ROI, n = 24: intima, n=6; media, n=9; adventitia, n=9), Non-stenosed (ROI, n = 27: intima, n=9; media, n=9; adventitia, n=9), and Stenosed (ROI, n = 29: intima, n=10; media, n=10; adventitia, n=9). **B.** Representative immunohistochemistry images of versican (red), aggrecan (red), and decorin (magenta) and their quantification (n=3, each group). Lumen area was marked as *. Scale bar = 100 or 200 μm.

To further examine ECM protein distribution, immunofluorescence staining was performed. Versican tended to be more abundant in the intima across all graft types and was significantly higher in the Non-stenosed intima compared with its media and adventitia (\*\**P*<0.01, **Fig. 5B**). Aggrecan was significantly enriched in the intima of Stenosed compared with both Denovo (\*\*\*\**P*<0.0001) and Non-stenosed (\*\*\**P*<0.001), and was also higher than in the Stenosed media and adventitia (\*\**P*<0.01, **Fig. 5B**). Decorin tended to be more abundant in the adventitia across graft types and was significantly higher in the adventitia of Denovo compared with its intima (\*\**P*<0.01, **Fig. 5B**).

### Cell Deconvolution Analysis and VSMC Phenotypic Feature

Cell deconvolution analysis was performed to better understand the composition of the graft samples. This approach used a single cell RNA sequencing dataset previously generated for the vasculature (https://cellxgene.cziscience.com). In the intima, endothelial cells (ECs) were significantly more abundant in Denovo compared with Non-stenosed and Stenosed (\**P*<0.05), whereas VSMCs were more prevalent in Non-stenosed and Stenosed than in Denovo (\**P*<0.05). In the media, ECs were likewise enriched in Denovo relative to Non-stenosed and Stenosed (\*\*\*\**P*<0.0001). In the adventitia, ECs were more abundant in Denovo (\**P*<0.05), while fibroblasts and macrophages were more prominent compared with the intima and media (**Fig. 6A**).

**Figure 6.**
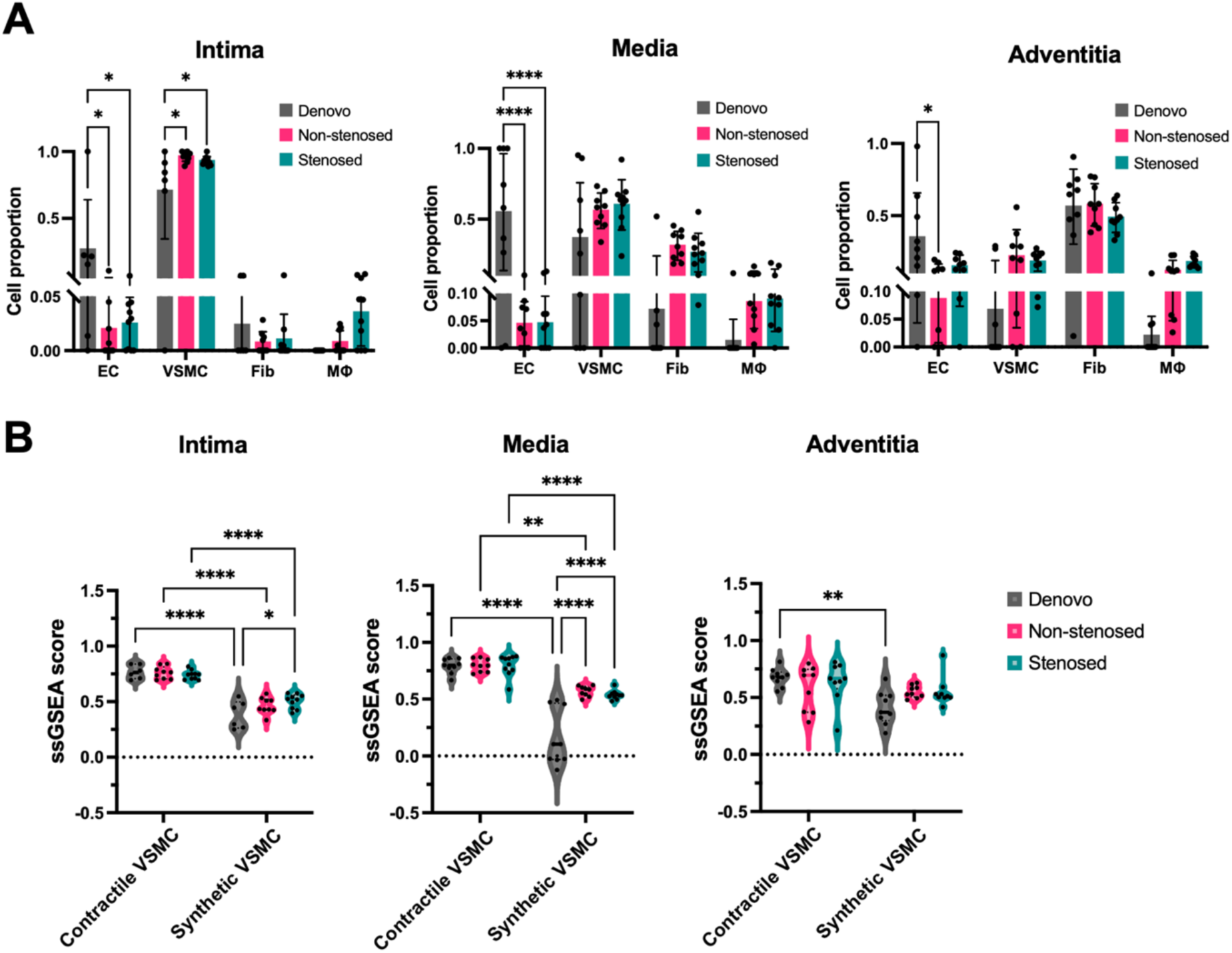
Cell deconvolution conducted using SpatialDecon R package and VSMC phenotypic features. **A.** Cell deconvolution analysis across graft layers. **B.** GSEA-based assessment of VSMC phenotypes, including contractile and synthetic subsets. Denovo (ROI, n = 24: intima, n=6; media, n=9; adventitia, n=9), Non-stenosed (ROI, n = 27: intima, n=9; media, n=9; adventitia, n=9), and Stenosed (ROI, n = 29: intima, n=10; media, n=10; adventitia, n=9).

To further assess VSMC phenotypic characteristics, ssGSEA was conducted. In the intima, contractile VSMCs were consistently more enriched across all graft types compared with synthetic VSMCs (\*\*\*\**P*<0.0001), although Stenosed grafts exhibited relatively higher levels of synthetic VSMCs than Denovo (\**P*<0.05). In the media, both Non-stenosed and Stenosed showed higher enrichment of synthetic VSMCs than Denovo (\*\*\*\**P*<0.0001). In the adventitia, although no significant differences were observed, Denovo grafts tended to exhibit higher levels of contractile VSMCs relative to synthetic VSMCs (\*\**P*<0.01, **Fig. 6B**).

## Discussion

In this study, we applied a spatial transcriptomic approach using DSP to human peripheral vein grafts to elucidate layer-specific molecular mechanisms underlying vein graft stenosis. By comparing de novo GSV, normally healed grafts, and stenosed grafts across vascular layers, we identified both similar and distinct transcriptional features associated with adaptive versus negative remodeling. Non-stenosed grafts were characterized by enrichment of pathways related to negative regulation of the cell cycle and proliferation, with increased expression of antiproliferative genes such as *CDKN1A* and *IGFBP3*. In contrast, stenosed grafts exhibited enhanced MAPK pathway activity, accompanied by reduced *DUSP1*-mediated regulation and a more inflammatory transcriptional environment. In addition, spatially distinct alterations in ECM remodeling—including differential expression of proteoglycans (*VCAN*, *ACAN*, *DCN*)—and a shift toward a synthetic VSMC phenotype were observed in stenosed regions. Overall, these findings are consistent with prior studies implicating coordinated regulation of cell cycle control, inflammation, extracellular matrix remodeling, and VSMC phenotype in vein graft remodeling, and suggest that spatially resolved analysis may provide additional insight into the molecular heterogeneity underlying graft adaptation and failure.

A characteristic of Non-stenosed grafts in our study was a transcriptional program indicative of regulated cell cycle control and restrained cellular proliferation. Non-stenosed grafts exhibited increased expression of *CDKN1A*, *LMNA*, and *IGFBP3* across vascular layers, with enrichment of GO terms related to negative regulation of cell population proliferation. *CDKN1A* was consistently enriched from the intima through the adventitia, and the MAPK regulator *DUSP1* was preferentially enriched in the media and adventitia compared with Denovo. In contrast, these antiproliferative signals were attenuated in Stenosed grafts, suggesting reduced protective cell cycle restraint during remodeling. Cell cycle dysregulation has been implicated as a driver of intimal hyperplasia and vein graft failure. Cyclin-dependent kinase inhibitors such as p21 (*CDKN1A*) and p27 (*CDKN1B*) limit vascular cell proliferation after injury, and reduced activity of these checkpoints has been linked to pathological neointimal expansion.^25, 26^ Genetic variation in p27 has also been associated with human vein graft failure and enhanced proliferation of venous adventitial cells, underscoring the importance of cell cycle control within the graft wall.^13, 27^ Another previous study has demonstrated that MAPK downstream signaling contributes to IH through activation of cyclic adenosine monophosphate response element binding protein via *MAPKAPK3* and *FHL5*, promoting VSMC proliferation and migration.^28^

These findings suggest that growth-inhibitory signaling is preserved across vascular layers in Non-stenosed grafts but attenuated in Stenosed grafts. These results further support the notion that impaired cell cycle control may contribute to maladaptive remodeling in peripheral vein graft stenosis.

Inflammatory regulation appears to play an important role in distinguishing adaptive from pathological vein graft remodeling, with the *DUSP1*–MAPK axis likely contributing to this process. *DUSP1* is a well-established negative regulator of MAPK signaling^29^ and has been shown in previous vascular studies to constrain inflammatory gene expression and limit VSMC proliferation following vascular injury.^30^ In addition, experimental studies using pharmacological MAPK inhibition have demonstrated that suppression of ERK1/2 activation attenuates medial cell proliferation, reduces inflammatory cell infiltration, and limits neointimal hyperplasia in vein graft models, supporting a causal role of MAPK signaling in graft remodeling.^31^ In the present study, *DUSP1* expression was higher in Non-stenosed grafts, particularly within the intima and media, suggesting the presence of a negative feedback mechanism that may dampen MAPK-driven inflammatory signaling during graft healing. In contrast, Stenosed grafts exhibited reduced *DUSP1* expression and increased Net MAPK activity, consistent with enhanced proinflammatory signaling. GO analysis further supported this pattern, as inflammation-related pathways were more frequently enriched in Non-stenosed grafts across vascular layers. Previous studies have suggested that inflammation is not inherently detrimental to vascular repair, but that its magnitude, duration, and resolution critically determine remodeling outcomes.^32, 33^ In line with this concept, macrophage analysis in our study revealed no significant differences in overall M1/M2 polarization between graft types. However, a modest enrichment of M1 macrophage signatures based on ssGSEA score was observed in the adventitia of Non-stenosed grafts compared with Denovo. The adventitia has been increasingly recognized as an immunologically active compartment that contributes to vascular remodeling through regulated inflammatory signaling.^34, 35^ Notably, this localized M1 enrichment in Non-stenosed grafts was accompanied by increased expression of anti-inflammatory regulators including *DUSP1* and *ZFP36*, which is known to promote decay of proinflammatory cytokine mRNAs.^36^ Together, these findings are consistent with a model of regulated or resolving inflammation during graft healing, whereas stenotic grafts may be associated with more persistent inflammatory activation. These observations suggest that vein graft outcomes may depend not only on the presence of inflammatory responses, but also on the ability to appropriately regulate and resolve them.^5^ ECM remodeling is an important component of vein graft adaptation after implantation and may exhibit layer-specific and graft type–dependent features. In this study, the large proteoglycans *VCAN* and *ACAN* were enriched in the intima of both Non-stenosed and Stenosed grafts, consistent with their established roles in neointimal formation and early responses to arterialization. Proteoglycans such as aggrecan have been implicated in vascular remodeling, including in stented coronary arteries where coordinated regulation of aggrecan and aggrecanases contributes to neointimal matrix reorganization.^37^ Similarly, differential regulation of versican across vascular layers has been reported in human vein grafts,^38^ supporting the concept that proteoglycan remodeling is spatially organized within the graft wall. The shared intimal upregulation observed in both graft types suggests that proteoglycan accumulation may represent a common adaptive response rather than a stenosis-specific feature. In contrast, the small proteoglycan *DCN* was preferentially enriched in the adventitia, supporting a potential layer-specific role in matrix organization. Decorin has been reported to regulate collagen fibrillogenesis and attenuate profibrotic signaling, and its adventitial predominance may reflect a stabilizing or reparative function during graft remodeling.^39–41^ Although differences between Non-stenosed and Stenosed grafts were modest, Stenosed grafts exhibited trends toward increased *COL1A1* expression and alterations in matrix turnover–related genes, including *MMP2* and *TIMP1*, which may be consistent with a shift toward collagen-dominant matrix remodeling. Gene expression patterns were not always fully concordant with immunohistochemical localization, likely reflecting post-transcriptional regulation, differential protein stability, and the prolonged extracellular persistence of matrix components.^42, 43^ Overall, these findings suggest that ECM remodeling in peripheral vein grafts is a spatially coordinated process, with subtle differences in matrix composition potentially contributing to the progression from adaptive remodeling to stenosis.

Cell-type composition and VSMC phenotypic features further characterized the remodeling landscape of the vein grafts. Cell deconvolution analysis indicated that ECs predominated across layers in Denovo, whereas both Non-stenosed and Stenosed grafts exhibited an increased proportion of VSMCs, particularly within the intima, consistent with structural adaptation following arterialization. This transition from an endothelial-rich venous state toward a VSMC-dominant architecture has been described as a hallmark of early vein graft remodeling and neointimal formation.^5^ Phenotypic analysis using ssGSEA demonstrated that contractile VSMC signatures generally remained more prominent than synthetic signatures across graft types and layers, suggesting partial preservation of differentiated characteristics. Nevertheless, modest differences were observed. In the intima, Stenosed grafts showed higher enrichment of synthetic VSMC signatures compared with Denovo, and in the media, both Non-stenosed and Stenosed grafts exhibited greater synthetic enrichment than Denovo. Although these differences were limited in magnitude, they are directionally consistent with prior studies demonstrating that VSMC phenotypic modulation—from a contractile to a synthetic state—is associated with intimal hyperplasia and graft remodeling.^44, 45^ Phenotypic switching of VSMCs has been linked to cell cycle activation, ECM production, and inflammatory signaling, suggesting integration of multiple remodeling pathways within the vessel wall. In this context, our spatial analyses suggest that subtle shift toward synthetic VSMC signatures, when considered alongside changes in cell cycle regulation and matrix remodeling, may be associated with progression toward stenotic remodeling. These findings support a role for VSMC phenotypic modulation in vein graft remodeling, although the effects were modest.

Several limitations of this study should be acknowledged. First, the sample size was limited (n = 3 patients), and all analyses were performed using FFPE specimens, which inherently restrict transcript coverage and sensitivity compared with fresh or frozen tissues. In addition, the analyzed graft samples predominantly represent an intermediate phase of vascular remodeling, as revision surgery was performed at approximately 10 months after the initial bypass. Therefore, molecular changes associated with early and late stages of graft remodeling were not captured in this study. Furthermore, although unused GSV segments were used as controls, previous studies have demonstrated that genes associated with IH, such as *MAPKAPK3* and *FHL5*, can already be expressed in pre-grafting vein segments, suggesting inherent biological variability among baseline samples.^28^ Second, the study design was cross-sectional, lacking temporal resolution.

Although immunohistochemical analyses provided complementary spatial information, functional validation using in vitro or in vivo models was not performed, limiting direct assessment of causal mechanisms. Third, cell deconvolution analysis relied on a publicly available single-cell reference dataset derived from general vascular tissues, as open-access single-cell RNA sequencing datasets specific to human veins or vein grafts are currently unavailable. As a result, the inferred cell-type proportions may not fully capture the precise cellular composition of vein graft tissues. In addition, the present spatial transcriptomic approach did not allow definitive identification of the specific cell types responsible for the observed expression of key regulatory genes such as *CDKN1A* and *DUSP1*. Future studies will aim to collect both FFPE and matched frozen specimens from the same vein grafts to generate dedicated single-cell RNA sequencing datasets. Integration of these datasets with spatial transcriptomic analyses is expected to improve the accuracy of cell deconvolution and enable more precise identification of cell-type–specific gene expression patterns. Despite these limitations, our results add pathophysiological significance to transcriptomic analyses of vein grafts. Another major strength of this study is the application of layer-resolved spatial transcriptomic analysis to human peripheral vein grafts, providing novel insights into the coordinated molecular programs underlying graft healing and stenosis. This approach offers a valuable framework for future mechanistic and translational investigations of vein graft disease.

In summary, this study demonstrates the potential of spatial transcriptomic analysis to elucidate mechanisms underlying peripheral vein graft stenosis using human specimens. Despite the limited sample size, our findings highlight coordinated, layer-specific patterns involving cell cycle regulation, inflammatory signaling, extracellular matrix remodeling, and VSMC phenotype that distinguish normally healed from stenotic grafts. These observations are consistent with prior experimental and clinical studies and support the concept that vein graft outcomes are associated with the balance of these interconnected remodeling processes. This study reveals layer-specific molecular heterogeneity not captured by bulk approaches and provides a framework for understanding graft adaptation and failure. These results may help inform future studies aimed at validating and targeting pathways that contribute to long-term graft patency.

### What are the Clinical Implications?

This study provides spatially resolved insight into molecular processes underlying peripheral vein graft remodeling in human patients. Our spatial transcriptomic analysis supports previous observations that graft remodeling involves coordinated regulation of cell cycle control, inflammatory signaling, extracellular matrix remodeling, and vascular smooth muscle cell phenotype across vascular layers. In particular, the observed balance between antiproliferative signaling and mitogen-activated protein kinase-associated inflammatory activity may represent a key determinant of adaptive versus maladaptive remodeling. Although preliminary, these results are consistent with prior experimental and clinical studies and support the potential value of targeting proliferative and inflammatory pathways to improve vein graft durability. Furthermore, spatial transcriptomic approaches may help identify layer-specific therapeutic targets that are not readily detectable by bulk tissue analyses. Future studies with larger cohorts and longitudinal sampling will be important to validate these findings and improve long-term graft patency.

## Non-standard Abbreviations and Acronyms

GSV: Great saphenous vein
IH: Intimal hyperplasia
VSMC: Vascular smooth muscle cell
ECM: Extracellular matrix
TGF-β: Transforming growth factor β
DSP: Digital Spatial Profiler
NBF: Neutral buffered formalin
ISH: In-situ hybridization
FFPE: Formalin-fixed paraffin-embedded
DEG: Differentially expressed gene
GO: Gene ontology
GSEA: Gene Set Enrichment Analysis
EC: Endothelial cell

## Acknowledgments

We thank Drs. Shreeram Akilesh, Kelly D. Smith, and Emily Beirne for their advice and support in performing GeoMx Digital Spatial Profiler. We also thank Drs. Scott R. Kennedy and Hong Xie for their help in performing RNA sequencing.

## Supplemental Material

Figure S1–S5

Major Resources Table

